# Principles of coarse-scale functional organization in occipitotemporal cortex

**DOI:** 10.64898/2026.02.18.706551

**Authors:** Laura M. Stoinski, Talia Konkle, Martin N. Hebart

## Abstract

Occipitotemporal cortex is known to process visually perceived objects, yet identifying general principles underlying its coarse-scale functional organization remains challenging for two reasons. First, previous work leaves open whether proposed organizational dimensions, such as animacy or real-world size, are useful for characterizing high-level vision and whether they generalize to more diverse, naturalistic stimuli. Second, object properties are often highly intercorrelated, making it difficult to determine which dimensions, if any, dominate coarse-scale occipitotemporal cortex organization. We addressed these challenges through detailed analyses of the cortical topography underlying 15 object properties, using densely-sampled fMRI responses to thousands of natural images, and focusing on animacy and size given their established prominence. Although our results confirmed many characteristics of the purported animacy-size organization, they revealed distinctions that challenge dominant notions of this organization, highlighting the importance of generalizing to large-scale naturalistic data. Beyond animacy and size, we found that many of the 15 properties could alternatively serve as organizational dimensions of occipitotemporal cortex, yet none emerged as clearly dominant across all OTC. Together, our findings reveal a more complex animacy-size organization than previously assumed and highlight challenges of assigning individual principles to account for OTC organization.

## Introduction

Our brain allows us to characterize and make sense of objects according to countless different criteria. This ability not only supports the recognition of the objects around us. It also enables us to determine, among others, whether things are alive or not, whether they are big or small, whether they can move or remain static, or whether they are hard or soft. Such distinctions help break down the complexity of the visual world for effective communication, flexible behavior, and efficient storage in long-term memory. Yet, how is it possible that we can so efficiently map the many different ways we can perceive objects to these many different facets that reflect our knowledge of the world? One possibility is that our visual system forms the foundation for an efficient mapping to behavior^1^, providing a basis set of mid-to-high-level features that structure our perception of the world. At a finer spatial scale, this organization accounts for category-selective clusters in high-level visual cortex that respond specifically to ecologically relevant categories such as faces, bodies, or scenes^2–4^. At a coarser scale, these features may underlie the overall topographic organization of high-level visual cortex^1,5–8^. Of note, this organization does not require committing to a high-level conceptual framework, where organizational principles are abstract conceptual distinctions that are directly computed by the visual system. Instead, these dimensions can be treated as descriptive axes that, at a coarser scale, align with these broad conceptual distinctions^9,10^.

Adopting this view of high-level visual cortex organization, we need to answer two questions to advance our understanding of its function: First, are there dominant axes of representational space that form the foundation for this mapping between vision on the one hand and semantics, memory, and behavior on the other? And second, are these dimensions related to specific conceptual properties, or are there, instead, many possible explanations that can account for the coarse-scale organization of high-level visual cortex?

Animacy and real-world size have been proposed as candidate fundamental dimensions of object representations for enabling the brain to efficiently characterize and categorize objects^8,11–13^. Neuroimaging studies have underscored their importance by demonstrating that these dimensions account for substantial variance in occipitotemporal cortex (OTC), organizing object responses into alternating gradients of animate-inanimate and large-small preferences (see Fig. 1c). Animacy and size have been proposed to map in a symmetric, tripartite fashion, with regions preferentially responsive to large objects, small objects, or animals regardless of their size^8^ (see also Fig. 1e).

**Fig 1.**
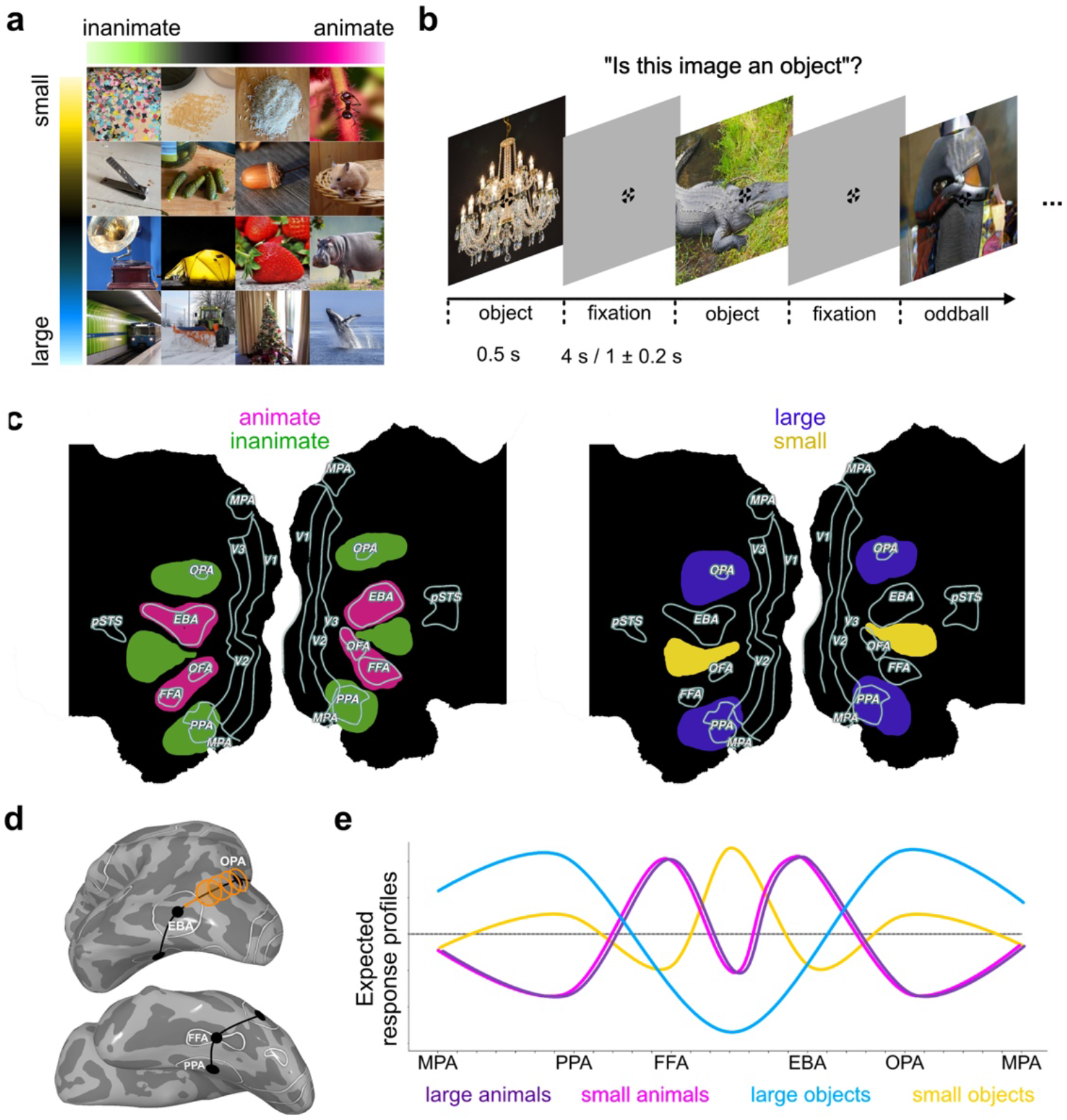
Methods and Hypotheses. **a)** The 8,640 images used in this study come from the THINGS object concept and image database^22^ and are associated with a diverse set of category ratings, including animacy and real-world size^23^. Displayed example images are similar to those used in the fMRI experiment but were taken from the public domain images dataset in THINGSplus^23^ for visualization purposes. **b)** Experimental procedure of the fMRI experiment. Participants viewed natural and synthetically generated object images while performing an oddball detection task. Only trials presenting natural object images were analyzed in the present study. **c)** Illustration of the expected animacy and size topographies^8^, dividing the cortex into alternating zones of animate-inanimate and large-small preference. **d)** Vector-of-ROI analyses. To visualize results as continuous response profiles, we defined a set of partially overlapping, spherical ROIs on a continuous path across the surface of high-level visual cortex. **e)** Illustration of the expected response profiles for large animals, small animals, large objects, and small objects across the spherical ROIs. Based on prior work^8^, we expected to find a tripartite organization, with regions showing distinct preferences for either large objects, small objects, or animals irrespective of their sizes.

Although animacy and size have been widely proposed as fundamental organizing principles of object representations, prior evidence largely stems from small-scale experiments using more controlled sets of object images^8,11–13^. However, much less is known about how these ideas generalize to the processing of more diverse and naturalistic stimuli, including images at the extremes of these properties, such as very large objects or less typical animals.

Beyond animacy and size, objects can be described by numerous, often intercorrelated properties. For example, animacy has been linked to properties such as moveability, agency, humanness, and curvature^13–19^, while size has been associated with manipulability vs. stability, and curvy vs. rectilinear distinctions^13,20,21^. Many previous studies did not focus on testing alternative organizational principles beyond animacy and size, or they typically investigated only a few properties at a time. To gain a more comprehensive understanding of object representations, it is therefore essential to elucidate the unique roles of animacy and size as organizational principles in relation to other object properties, whether other dimensions would better account for this organization, or whether coarse-scale responses in high-level visual cortex would be difficult to reduce to particular high-level conceptual properties. However, given the high correlation among known properties, teasing apart the effects of distinct properties has remained challenging.

In this project, we aimed to identify whether there are fundamental organizational principles that may dominate the large-scale functional organization of the occipitotemporal cortex, and if so, to what degree these dimensions correspond to specific high-level object properties. We focused our investigations on the well-known examples of animacy and real-world size, given their purported importance as highlighted in the literature^1,8,11–13^. First, we tested the extent to which animacy and size generalize as organizational principles to the processing of more diverse and naturalistic object images. Second, we examined the proposed importance of animacy and size in high-level visual processing relative to other object properties. Specifically, we tested whether 13 related properties could serve as alternative organizational dimensions by assessing how well they organize OTC responses into smooth topographical maps and by comparing their explanatory power for high-level visual response patterns. If animacy and size are truly fundamental principles, they should underlie the smoothest topographical organization and stand out clearly from the other properties in explaining OTC responses. If, however, multiple properties could serve as organizational principles, or other properties emerge as more dominant, this would suggest that our evidence favors no particular property, raising the question of whether comparing individual properties is the most informative approach for advancing our understanding of representational structure in high-level visual cortex.

## Results

First, we asked to what degree previously proposed topographic organizational principles in occipitotemporal cortex with respect to animacy and size^8,13^ generalize to a more diverse and naturalistic set of object images. To this end, we used the THINGS-fMRI dataset^22,24^ consisting of three densely sampled participants who were presented with images of 720 object categories of continuous animacy and real-world size ratings provided as part of THINGSplus^23^. This allowed us to evaluate the degree to which the previously proposed macro organization by animacy and size^8^ generalizes to more representative object images of more continuous variations in these properties. We then conducted a voxel-wise regression analysis using z-scored animacy and size predictors. The predicted pattern of results is highlighted in Fig. 1c.

### High generalizability of animacy organization, more pronounced partitioning of size organization

Fig. 2 illustrates the resulting beta coefficients of animacy and size, mapped onto the flattened visual cortex. Across all three participants, we replicated the proposed large-scale animacy preference map^8,13^, which divides OTC into alternating zones of inanimate and animate preferences (Fig. 2, left). Known category-selective regions mapped smoothly onto this animacy organization, with the face- and body-selective regions fusiform face area (FFA), occipital face area (OFA), and extrastriate body area (EBA) exhibiting a preference for animate categories, and the scene-selective occipital place area (OPA) and parahippocampal place area (PPA) responding more strongly to inanimate categories. Together, these results confirm the previously reported animacy organization and extend it to a much broader naturalistic dataset^24^.

**Fig 2.**
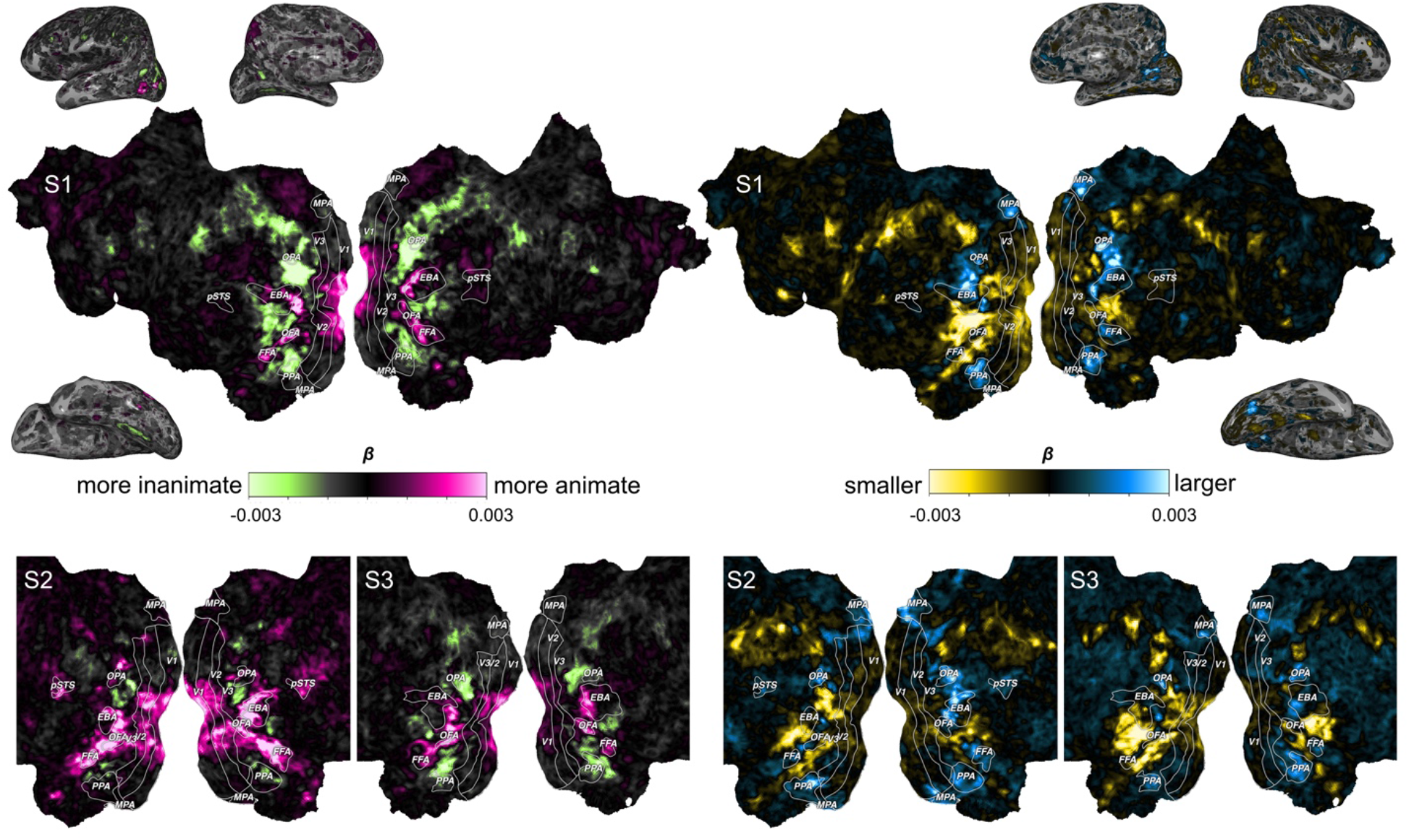
Animacy and Size topographies. Voxel-wise regression coefficients of the animacy (left) and size (right) predictors mapped on the flattened cortex. Color scales indicate the polarity and amplitude of unstandardized regression coefficients (*β*). The results mirror the characteristic alternation of animate to inanimate and large to small gradients^8,13^. However, beyond prior findings, we found an additional, bilateral small-preference cluster between FFA and PPA and pronounced size preferences within FFA, consistently in all three participants.

Regarding the size organization, we found a similar topography as in previous studies^8,13^, with a preference for large objects in PPA and OPA and a preference for smaller objects between FFA and EBA (Fig. 2, right). However, compared to previous work, the size organization was more asymmetric and partitioned, including a preference for larger objects in FFA and an additional preference zone for smaller objects between FFA and PPA. In participants 1 and 3, we further observed preferences for smaller object categories in OFA of the right hemisphere, with weaker yet qualitatively similar effects in participant 2. Thus, while these results partially replicate previous findings, they highlight a more fine-grained response profile for real-world size than reported previously.

We conducted various exploratory analyses to evaluate the robustness of these findings. These included analyses in volumetric data instead of surface vertices, probing preferences for the 25% largest and smallest objects rather than a continuous size range, restricting the stimulus set to objects within a comparable size range or the same object categories as in the original studies^8,13^, and testing the effects of object eccentricity, object size in the image, and fine-grained clutter in images with small objects (see Supplementary Methods and Supplementary Fig. 1 for details on these analyses). The reported results remained robust to these variations, highlighting the stability of these findings in relation to real-world size.

### Representation of all four quadrants in the animacy-size organization

Having characterized the coarse-scale topographic organization of animacy and real-world size in our dataset, we next determined the interaction of both properties.

Previous work has suggested a tripartite organization^8^, with a distinction between large objects, small objects, and animals irrespective of size (Fig. 1e). To align more closely with the analyses in previous work, we collapsed the continuous ratings to binary categories and selected only a subset of the 720 objects, focusing on the 25% largest, smallest, most animate, and most inanimate objects (Fig. 3a). Next, we conducted a vector-of-ROI analysis (Fig. 1d), defining a set of partially overlapping spherical ROIs that mapped a continuous path across high-level visual cortex. We then computed the mean response for each group within each spherical ROI, normalizing the values by subtracting the average response across all four groups within the respective ROI.

**Fig 3.**
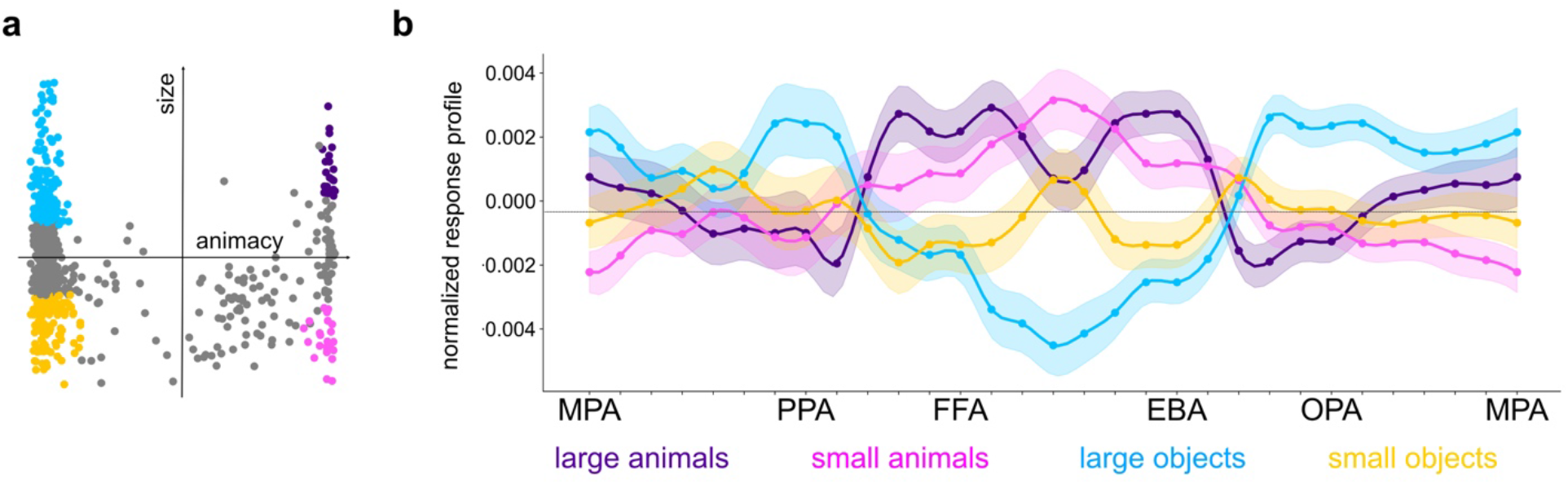
Relative response profiles to large and small animals and objects. **a)** Objects plotted in the animacy-size space, colored by their group membership. Animals included only non-human animals and objects included only very inanimate categories. Large and small classifications were defined using the upper and lower 25th quartiles of the largest and smallest animals or objects. Only colored objects were considered in the comparisons of response profiles (*n* = 308). **b)** Response profiles to large and small animals and objects across spherical ROIs. The group responses were normalized using the grand mean across groups. The errors represent the SEM*1.96 across members of each of the four groups. While our results coarsely align with those found by Konkle & Caramazza^8^ (see also expected profiles in Fig. 1e), we observed more pronounced differences between large and small animals than previously reported, as well as generally stronger responses to animals compared to small objects in the previously proposed small-object preference region between the FFA and EBA.

Matching the results of Konkle and Caramazza^8^, we found peak preferences for large objects in PPA and OPA, animals in FFA and EBA, and a stronger preference for smaller than larger object categories between FFA and EBA (Fig. 3b). However, in contrast to previous work that found no difference between large and small animals (Fig. 1e), our results exhibited a notable preference for larger animals over smaller animals in FFA and EBA. Furthermore, while previous research identified a peak preference for small inanimate objects in the region between the FFA and EBA, we found that this region showed the strongest responses to small animate categories. The objects eliciting the highest or lowest response in spherical ROIs corresponding to pronounced animacy or size preferences can be found in Supplementary Table 1.

We tested whether the larger differences within the animal domain remained robust when considering only the categories included in the original studies. The results remained consistent, suggesting that these findings are not a consequence of our more diverse set of animals. Thus, our results reveal a more complex interaction between animacy and size than suggested by the tripartite model, indicating that size may serve as an organizational dimension within both the inanimate and animate domains, allowing for a complete separation of objects and animals across sizes. This highlights that high-level visual cortex already allows categorization not only according to individual object features, but with respect to meaningful differences between large and small animate and inanimate objects.

### Assessing the role of diverse alternative principles in providing fundamental dimensions of high-level visual cortex organization

While the coarse-scale organization of high-level visual cortex has been hypothesized to be governed by animacy and real-world size^8^, little is known about the relative importance of these properties with respect to other candidate principles for a broad set of natural object images. This uncertainty stems from the high collinearity of many other object properties and from the limited scale of previous studies that have attempted to disentangle animacy and size from related dimensions^8,13,14,16–18,21^.

To address this issue, we tested the purported importance of animacy and size as fundamental organizational principles of high-level visual cortex with respect to other candidate principles for a broad set of natural object images. We reasoned that a good organizational principle should fulfill two criteria: First, produce smooth, coarse-scale topographic maps, and second, explain considerable variance across high-level visual cortex. Assuming that our aim was to reduce the organization of high-level visual cortex to only a handful of properties, we tested whether animacy and size were among the best-suited candidates. Disentangling animacy and size from these highly collinear variables is only possible given the large size of our dataset, featuring object images with many diverse combinations of these properties.

To clarify the relationships among object properties, we first computed pairwise Pearson correlations (Fig. 4a) and visualized their relationships using hierarchical clustering analysis with average linkage (Fig. 4b). Animacy ratings were most strongly related to naturalness, agency, and the ability to move. Real-world size ratings showed the strongest positive correlation with heaviness and hardness, and the strongest negative correlation with the ability to be moved and manipulability.

**Fig 4.**
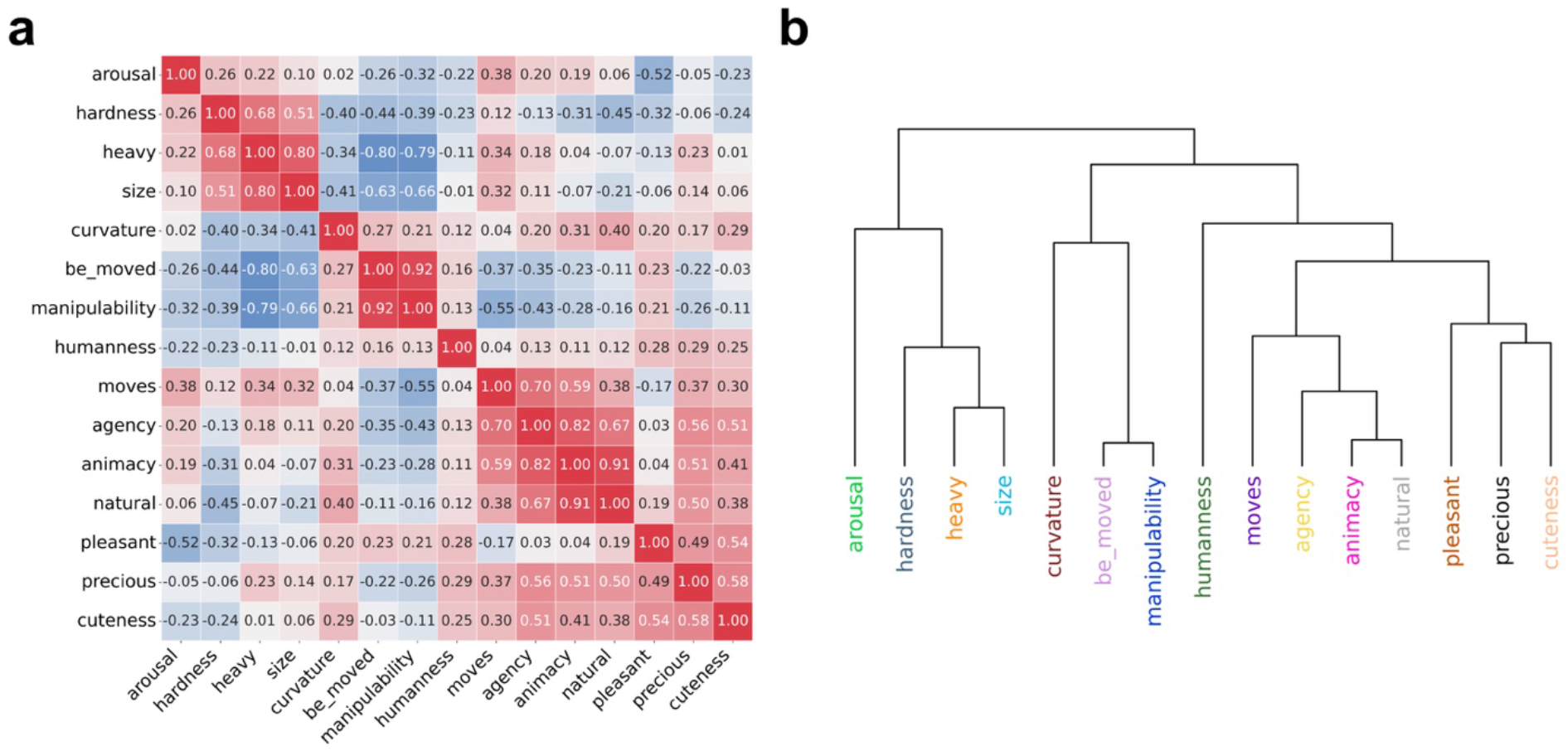
Correlations and hierarchical clustering of object properties. **a)** Correlation matrix of the 15 object properties. Several properties were strongly intercorrelated. Animacy showed the strongest positive correlations with naturalness and agency. Size was most strongly positively correlated with heaviness and most strongly negatively correlated with manipulability and ability to be moved. **b)** Hierarchical clustering dendrogram of object properties. The dendrogram illustrates how properties cluster at different levels of similarity.

As the first test of the importance of animacy and size, we evaluated to what degree the 15 different properties would organize occipitotemporal cortex responses into smooth coarse-scale gradients. If there were special roles of animacy and size, we would expect them to clearly stick out from the other properties when it comes to organizing OTC responses into smooth and broad maps. In contrast, if other properties produce similarly smooth maps, this would suggest that they qualify as alternative organizational principles, challenging the purported importance of animacy and size as primary dimensions of representational space.

We performed voxel-wise univariate OLS regression analysis, showing that all of our properties organize OTC responses into broad topographical maps. As a control, shuffling property ratings (e.g. size) resulted in a substantially less smooth topography. The spatial organization of properties indeed mirrored their correlation structure, with properties strongly correlated with animacy, such as moveability, agency, humanness, naturalness, and preciousness, showing topographical organizations similar to the animacy organization. Likewise, properties correlated with size, such as manipulability, curvature, heaviness, and the ability to be moved, showed a topography resembling the size organization (Fig. 5). Thus, animacy and size do not distinguish themselves in their ability to produce broad, smooth topographical maps, and other properties qualify as alternative candidate organizational principles.

**Fig 5.**
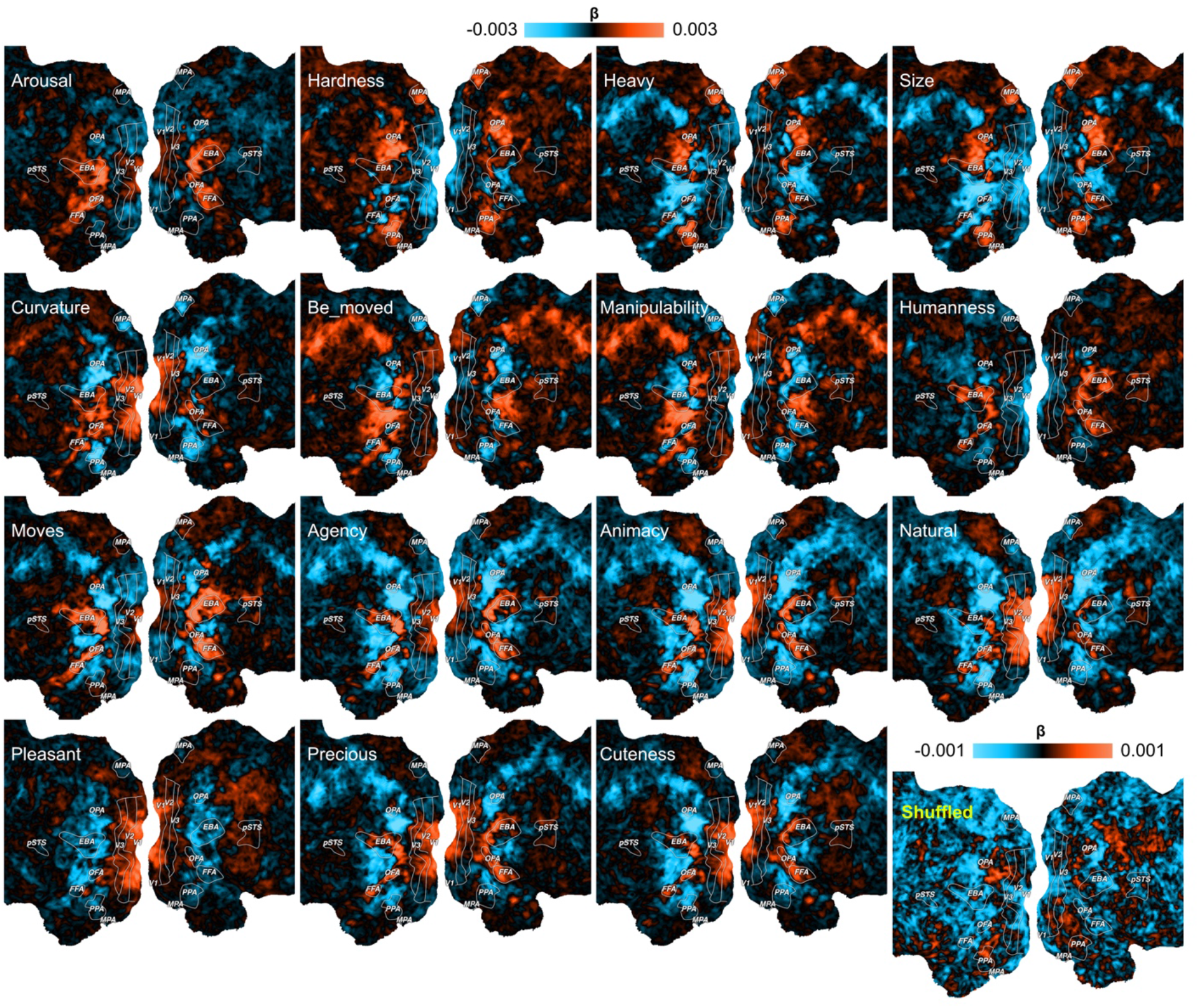
Topographic maps of object properties. Univariate regression coefficients (*β*) of object properties and shuffled ratings (control condition) mapped on the cortex of one example participant, S1. The regression coefficients for the shuffled ratings were smaller in magnitude. Thus, a different color scale was used to enhance the visibility of their topographic pattern. All properties reveal a smooth coarse-scale organization of high-level visual cortex, with several resembling the organization observed for animacy or real-world size. In contrast, maps based on shuffled size ratings did not show smooth gradients, indicating that the observed topographies for object properties reflect meaningful neural structure rather than noise.

To quantify the smoothness of each property coefficient map (Fig. 5) more objectively, we computed a neighborhood-based total-variation score on the cortical surface, weighting absolute differences in regression coefficients between vertex pairs by their geodesic distance, with lower scores indicating smoother maps (Fig. 6, top). We also report a variant of the score that further weights the absolute differences in regression coefficients by the vertices’ reliability (derived from THINGS-fMRI^24^; Fig. 6, bottom). The results reflect what was expected from the property topographies and the correlational structure among properties. All 15 tested object properties showed smoother maps than a shuffled property baseline, indicating that all of them fulfilled the criterion of a coarse-scale gradient. While there was some degree of variability in the scores, animacy and size did not specifically stick out with respect to the others.

**Fig 6.**
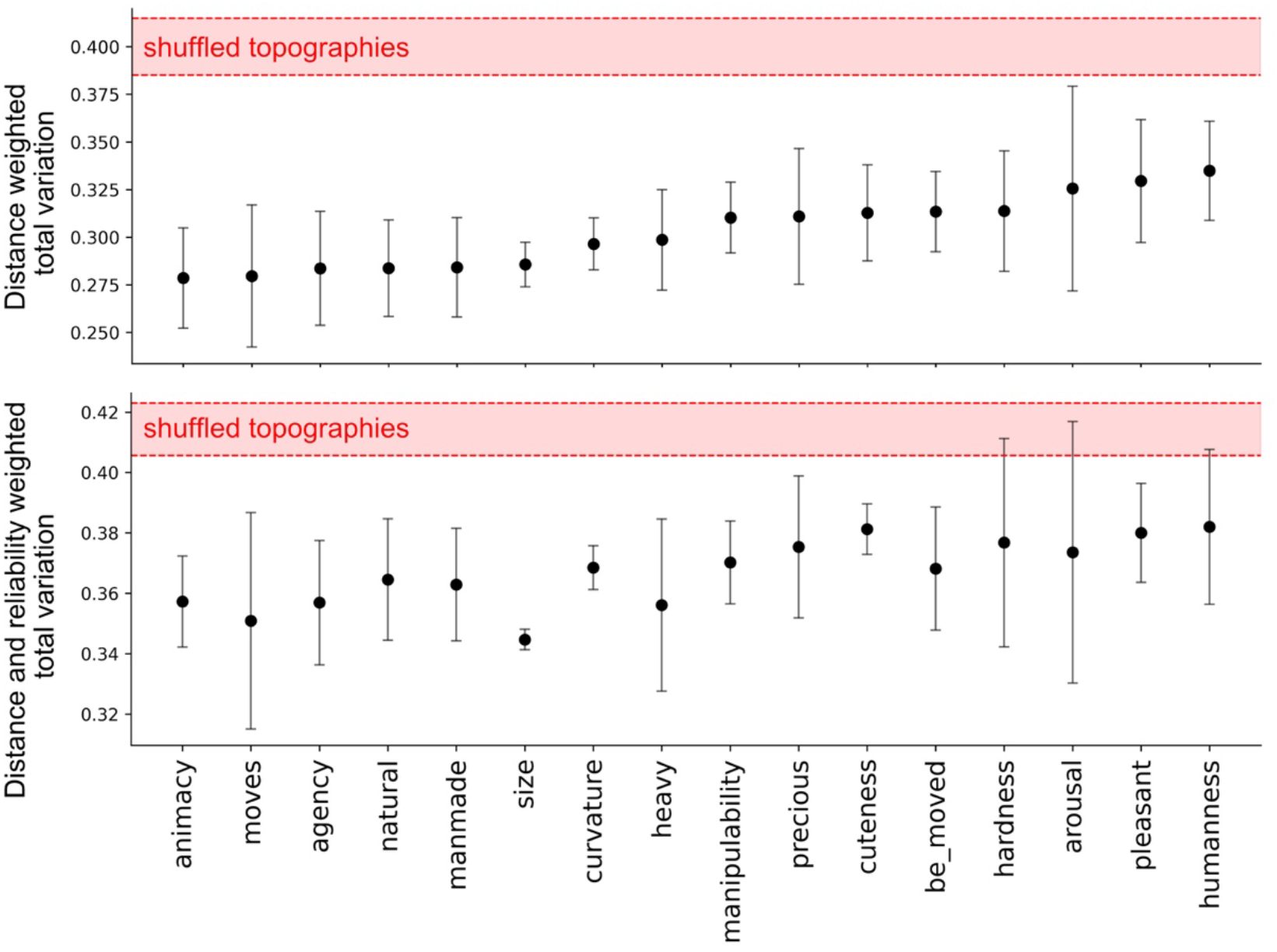
Quantification of topographical smoothness. Distance-weighted and distance-reliability-weighted total variation of cortical property coefficient maps, averaged across participants. Lower scores indicate smoother maps. Error bars show the bootstrap confidence interval derived from 1,000 resampled topographies. Many properties show comparable smoothness, with neither animacy nor real-world size uniquely standing out.

Properties that correlated with animacy or size appeared to show similar smoothness, although this was not always the case. For example, naturalness was highly correlated with animacy but was less smooth than moveability, a less strongly correlated property. To test whether a property’s smoothness merely reflects its correlation with animacy, we regressed animacy out of the property scores and fMRI responses. After removing the variance shared with animacy, many properties continued to produce broad and smooth topographical maps (Supplementary Fig. 2). This suggests that their smoothness cannot be explained solely by their association with animacy.

Together, these results show that several properties produce similar broad and smooth functional topographies and thus do not support a unique role of animacy or size in higher-level visual cortex organization.

As a second test of whether animacy and size are among the best-suited organizational principles of high-level visual cortex organization, we disentangled their unique contributions in explaining high-level visual response patterns compared to the other properties.

To provide a baseline of how much variance is captured by all 15 object properties conjointly, we first ran a full model incorporating all properties. The full model predominantly accounted for variance in high-level visual cortex and captured large fractions of explainable variance (Fig. 7a), highlighting that the 15 object properties indeed reflected high-level visual responses well.

**Fig 7.**
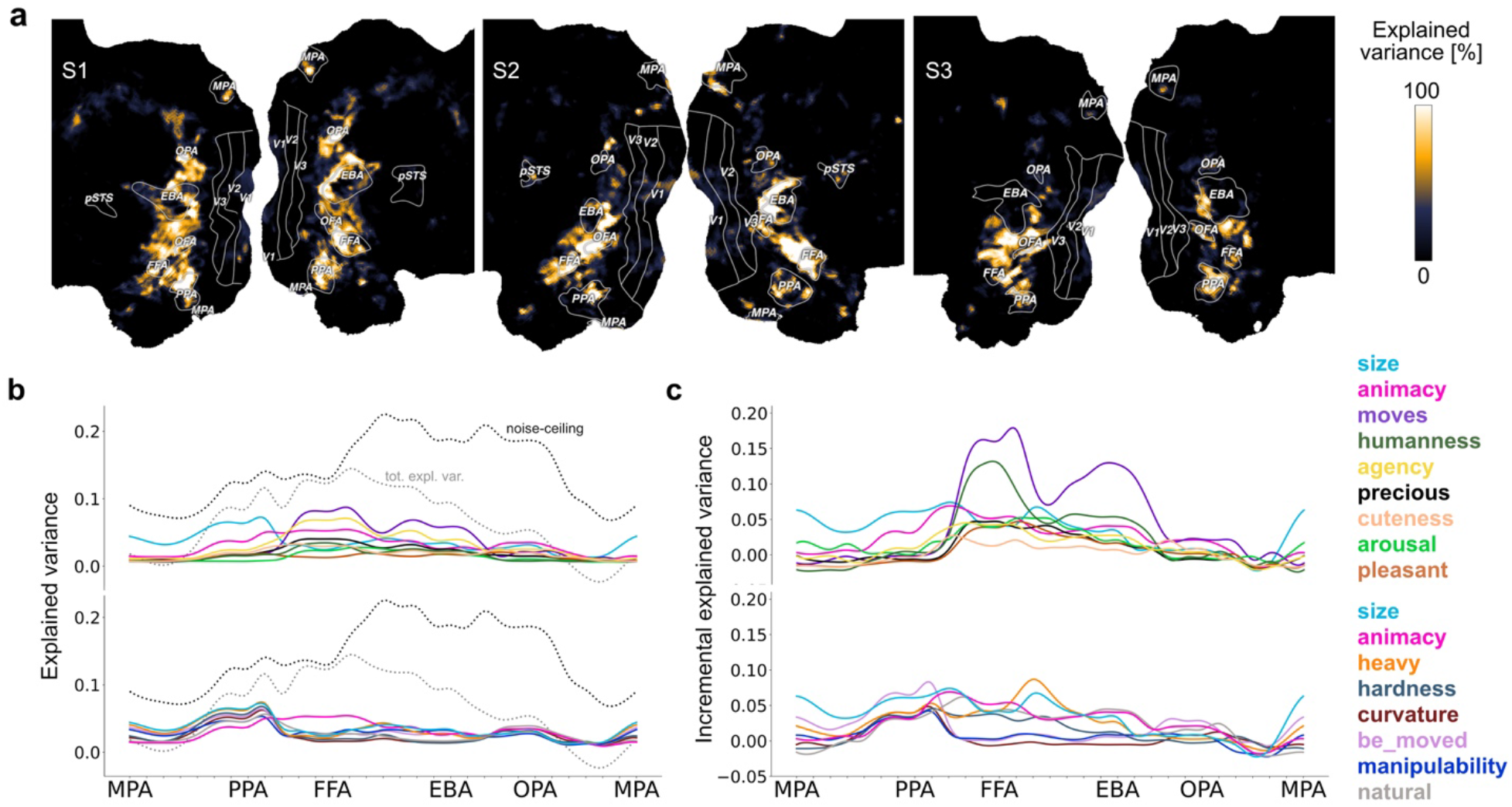
Results of the hierarchical partitioning analysis of object properties. **a)** Variance explained by an encoding model including all fifteen properties (cross-validated and noise-normalized) mapped on the flattened cortex of each participant. **b)** Explained variances of univariate models for each property (cross-validated) across spherical ROIs and averaged over participants. Response profiles are divided into two graphs for improved visibility, with animacy and size featured in both. Before disentangling the properties from each other, several properties accounted for variance comparable to or greater than that explained by animacy and size across high-level visual cortex. Note that these results are not noise-normalized, allowing the noise ceiling to be displayed as a reference. **c)** Results of the hierarchical partitioning. Incremental explained variances of each property (cross-validated and noise-normalized) across spherical ROIs and averaged over participants. Response profiles are divided into two graphs for improved visibility, with animacy and size featured in both. After disentangling them from the other properties, animacy and size still accounted for incremental variance across the high-level visual cortex. However, other properties explained similar or even larger shares of variance in regions previously associated with animacy or size preferences.

As a second baseline of how much variance each property explained across high-level visual cortex without disentangling it from the others, we computed a univariate regression for each property (Fig. 7b). All properties captured variance in high-level visual regions, with moveability and agency explaining the largest amount of variance in regions previously associated with animacy preferences, and size and heaviness explaining the most in regions previously related to size preferences. Supplementary Tables 2 and 3 provide additional information about which objects were best and worst predicted by the individual property models in FFA and EBA, where the most pronounced effects were observed.

Finally, we disentangled the contribution of each of the 15 properties in explaining high-level visual cortex responses by determining the incremental variance explained by each property. While variance partitioning has been used in the past to determine unique and shared variance in explaining patterns of brain activity for collinear variables^25^, unique variance is no longer a good indicator of added value of a specific variable when dealing with a larger number of collinear variables, since highly interrelated variables would share a large fraction of variance to start with and thus would have a smaller capacity for explaining unique variance. To reduce this challenge and still report the unique added value of each individual feature, we resorted to an alternative analysis method known as hierarchical partitioning, which provides a more accurate assessment of the independent contribution of each individual variable^26^. This method involved computing the incremental explained variance when a property was included in a model, compared to the identical model without that property, averaged across all possible 2^14^ model combinations per property.

The incremental contribution of each property in explaining high-level visual cortex responses, averaged across participants, is shown in Fig. 7c. Results at the individual participant level are shown in Supplementary Fig. 3. Many properties accounted for sizable incremental variance, but none consistently dominated across all regions. Animacy accounted for incremental variance in several regions of high-level visual cortex. However, moveability explained considerably more incremental variance in FFA and EBA, regions which previously have been associated with animacy preferences^8,13^, while humanness contributed considerable additional variance in FFA. After disentangling the role of size from the other properties, size continued to account for considerable incremental variance across high-level visual cortex. However, the ability to be moved explained slightly more incremental variance than size in PPA and OPA, regions previously linked to large-object preferences^8,13^. Meanwhile, size and heaviness explained the majority of variance in the region between the FFA and EBA, which was previously associated with small-object preferences^8,13^.

To determine whether the high collinearity between animacy and naturalness, or between size and heaviness, contributed to the smaller-than-expected effects of animacy and size, we re-ran the hierarchical partitioning analysis without naturalness and heaviness (Supplementary Fig. 4a). The results were highly similar to those of the full model including all 15 properties, with moveability and humanness continuing to explain the most incremental variance in FFA and EBA. We additionally ran a model without naturalness, heaviness, and agency to further reduce the number of predictors that were strongly correlated with animacy (Supplementary Fig. 4b). In this model, moveability and humanness still explained the largest amounts of incremental variance, although the incremental variance explained by animacy in EBA was now similar to that explained by humanness. Together, these exploratory analyses suggest that the pronounced effects of moveability and humanness were not driven by excessive collinearity among animacy and other predictors.

In summary, all investigated properties contributed to variance in high-level visual cortex, but none dominated in explaining OTC responses across all regions. If a single property were to be prioritized, then moveability may be a stronger organizational principle than animacy, however its effects appeared concentrated in FFA and EBA.

## Discussion

Is there a basis set of dimensions that act as fundamental organizational principles of high-level visual cortex, allowing our visual system to efficiently map visual input to our semantic knowledge of the world? If so, which properties or features dominate the functional organization? Previous work has proposed a parsimonious framework for the organization of high-level visual cortex with respect to the properties of animacy and real-world size. Animals, small objects, and large objects are often linked to different functions such as social interaction, manipulation, and navigation^1,8,12,13^. From this perspective, animacy and size were proposed to act as highly informative properties that allow us to efficiently map visual information to long-range networks tuned to distinct behavioral demands^1,27^.

Here, we aimed to provide a more comprehensive test of whether animacy and size clearly stand out in providing fundamental axes of high-level visual organization or whether there are other properties offering an alternative account of high-level visual cortex organization. First, we tested the extent to which previous findings^8,13^ on the animacy-size organization generalize to more diverse and naturalistic object images.

Second, we evaluated the relative importance of 15 candidate object properties, including animacy and real-world size, by testing how smoothly they organize high-level visual cortex into coarse-scale gradients and by assessing the incremental contributions to explaining response patterns across the occipitotemporal cortex compared to 13 other candidate properties.

Our results cast previous findings of the animacy and size organization in a new light. First, our results confirmed the large-scale organization of animacy and extended it to a much larger range of natural object images^24^. For real-world size, we found a more asymmetrical and partitioned size organization than reported previously^8,13^. This result is in line with findings in another dataset that showed preferences for large objects near FFA and a preference for small objects dorsolaterally to PPA^28^. This suggests that the previously reported size organization may not fully generalize to responses to more naturalistic and diverse object images. While we explored several potential explanations for the more partitioned size organization, we could not identify the reason for the divergence in findings. Future studies are needed to directly replicate the original findings of the real-world size organization and test what differences lead to changes in response profiles across high-level visual cortex.

Second, we observed greater response differences between large and small animals than in previous studies^8^. Even when we limited our analysis to the categories featured in the original study^8^, the results remained broadly consistent. Other research utilizing naturalistic stimuli has also demonstrated response variations for animals of different sizes^29^. Together with our results, this suggests that, contrary to what has been shown in prior studies, the processing of both inanimate and animate objects can vary depending on their size, revealing a largely mirror-symmetric structure that does not fit a tripartite model. We further show that when using naturalistic objects and a large-scale dataset, the functional organization of the OTC differs from patterns observed in smaller, more controlled datasets.

Having identified the animacy and size organization, we tested their relative importance as possible organizational principles of high-level visual cortex. Building on important prior efforts to decorrelate animacy and size from other variables^13–18,21^, we made use of a comprehensive dataset of thirteen perceived object properties^23^ to examine their relationships with animacy and size simultaneously.

We demonstrated strong intercorrelation of object properties and showed that several properties performed similarly well at describing the organization of high-level visual cortex responses through broad and smooth gradients. This suggests that animacy and size are not unique descriptors of coarse-scale organization in high-level visual cortex. Rather, other object properties provide alternative accounts of a very similar large-scale representational structure.

When disentangling the incremental contribution of each property using hierarchical partitioning, we found that many variables explained incremental variances in distinct regions, with none dominating in all regions. If one were to pick, moveability accounted for more incremental variance than any other property, however, only in regions overlapping with or surrounding FFA and EBA. This finding is consistent with prior work from smaller, more controlled studies, which have already pointed out the limitations of animacy as strictly defined by the dichotomous distinction of animate and inanimate^14,16,18,19,30,31^.

Do these results imply that we should move away from the dominant notions of animacy and size, and instead, focus on moveability? Our results demonstrate that several properties explained notable variance across high-level visual cortex, while none clearly dominated across all regions, with a similar pattern of results found for incremental variance. Thus, even though it is generally appealing to identify simple laws that underlie complex observations, our results indicate that the organization of high-level visual cortex may be more complex than can be captured by a small number of conceptual properties.

Given the strong focus of our study around the properties of animacy and size, it can be argued that we tested a non-representative set of conceptual properties, and that there may be a different, small set of other properties that better capture responses in high-level visual cortex. At the same time, our study included many properties highlighted in previous research^1,13,16,17^ and other data-driven work has shown that early components show some correspondence with animacy^5,32–34^. Therefore, it is unlikely that other untested, vastly different conceptual properties would dominate responses in high-level visual cortex. Still, it remains possible that our analyses missed such properties and that high-level visual cortex is dominated by a small number of semantic axes. Aside from curvature, we also did not examine visual properties more directly, such as material properties^35^ or shapes^5^, which have been suggested to play a role in visual cortex organization. In addition, our approaches were purely linear, making it possible that non-linear mappings exist between individual properties and high-level visual cortex that capture much more variance.

Another possibility is that dominant axes exist, but that they do not map neatly onto the semantics we used to describe the world with our language. For instance, they may reflect mixtures or interactions of several properties^33,34^, or experience-based sensitivities for visual features that covary with conceptual properties^10,13,36,37^, which would indicate that high-level visual cortex may not easily correspond to individual conceptual properties.

The view that there may not be a clean mapping onto a small number of dominant semantic axes while additional axes still explain additional unique variance would dovetail nicely with the view that OTC may provide the functional basis for a highly diverse read-out of many object properties. This view provides a re-emphasis of high-level visual cortex function with the downstream functions of vision^1,6,38^, including memory, language, and decision-making, instead of being organized primarily along any particular axis. Importantly, these views need not be mutually exclusive. High-level visual cortex organization could be the outcome of multiple constraints imposed by both the statistical structure of the visual input and the optimized read-out of the diverse behavioral demands the visual system must support^1,38^.

One limitation of our work is that we did not investigate potential interactions beyond animacy and size, as testing these would require an unfeasibly large number of comparisons across all properties. Future research may selectively investigate the relationship between pairs or triplets of properties.

Second, object properties were assessed at the object level and may therefore not fully reflect how participants perceived the individual images shown during the fMRI experiment. However, the analyses were conducted on fMRI responses averaged across all 12 images per object, reducing the influence of such image-specific variation.

Further, as we conceptualize the fundamental organizational principles as continuous dimensions, we did not disentangle the contributions of moveability and humanness from category-selective responses to faces or bodies. However, while category selectivity may offer an alternative characterization of some aspects of high-level visual cortex organization^39^, it would only provide an incomplete picture of its coarse-scale functional organization. Given that much of the variance in high-level visual cortex was captured by the 15 properties, these likely included much of the variance captured by category selectivity. While category selectivity may thus include the broad representational goal of high-level visual cortex^40^, future work may specifically test the relative role of category selectivity in the high-level description of visual cortex.

Finally, it is possible that a purely data-driven approach could have yielded interpretable latent factors directly from the fMRI data. However, the single-trial reliability in the present data did not allow for such analyses, and similarly to above, given the explainability of the existing variables, these factors would likely reflect factors related to our 15 properties or their combinations. Future work may determine latent factors underlying neural object similarities and the properties and features they most closely capture.

In conclusion, our study broadens the scope of the previously suggested organization of high-level visual cortex with respect to animacy and real-world size. First, we revealed a more complex animacy-size organization than previously proposed. Second, we showed that other properties compete with, and often explain variance beyond, animacy and size. Thus, when aiming at understanding the nature of visual representations across high-level visual cortex, our findings underscore the importance of using diverse, naturalistic datasets for drawing more generalizable conclusions, while also raising the question of whether searching for individual organizational properties is sufficient to advance our understanding of its representational structure.

## Methods

### Ethics statement

The experiment was approved by the Ethics Committee of the Medical Faculty of Leipzig University (157/20-ek).

### Experimental design

#### Participants

Most object properties were drawn from the existing norm database THINGSplus^23^. Additionally, we collected a novel dataset of humanness, agency, cuteness, and hardness ratings. Participants for both existing and new ratings were recruited on the online crowdsourcing platform Amazon Mechanical Turk (AMT).

Novel property ratings were collected in separate tasks, one for each property. We recruited 3,307 unique workers from the United States who could participate in all kinds of tasks and in as many small sets of each task (HITs) as they liked. Participants provided informed consent in compliance with the Ethics Committee of the Medical Faculty of Leipzig University, Germany, and received small compensation for each completed HIT. We flagged potential carelessness if a participant completed ten trials of a HIT in less than 1.5 seconds, five trials in less than 1 second, or had a response variance of one or less across all trials, indicating repetitive button presses. If a participant was marked as insufficiently careful in three or more HITs, all their HITs were removed from data analysis, and they were prevented from further participation. After exclusions, 2,246 workers remained, completing a total of 23,814 HITs. Among them, 1,199 identified as female, 1,041 as male, and six as other gender, with a mean age of 37.43 (SD = 10.71).

We followed a similar procedure for the existing property ratings as described above. Details on participants and exclusion criteria are reported in Stoinski et al.^23^. Brain data were drawn from THINGS-data^24^, which includes fMRI responses from three healthy participants (two female, one male, mean age = 25.33 years).

#### Object properties

The novel dataset of perceived humanness, agency, cuteness, and hardness was collected in separate tasks. For each property, participants were asked to rate all 1,854 objects of the THINGS database^22^ on a scale from 1 to 9, asking, “How similar is this object to a human?” “Can the object act independently?” “How cute is this object?” and “How hard is this object?” Each object concept was sampled 40 times. Due to insufficient split-half reliability, an additional 40 samples per concept were collected for the humanness task.

All other property ratings were drawn from the THINGSplus database^23^. Size ratings were collected in two steps. In the first step, participants were presented with an object noun and asked to indicate the approximate size on a continuous scale with 520 units. As reference points, we aligned nine objects along the scale (i.e., ranging from “grain of sand” through “microwave” to “aircraft carrier”). In the second step, the rating scale zoomed in on the chosen interval, and one additional reference object was embedded between the initial reference objects. Some object categories may vary in size (e.g., some boats are much larger than others). To consider this variability, we asked participants to refine their initial answer by indicating the size range, capturing the smallest to the largest size of the object in the real world.

Animacy, naturalness, preciousness, ability to move, ability to be moved, heaviness, pleasantness, arousal, manmadeness, holdability, and graspability (with the latter two later combined into manipulability) were gathered in a single survey.

Participants employed 7-point Likert scales to assess how well each property described a given object noun (for a detailed description of data collection and validation, see Stoinski et al.^23^). Curvature was collected in a separate study in which participants rated the relative balance of curved versus rectilinear object contours in individual images on a scale from 0 (very rectilinear) to 100 (very curvy).

#### fMRI dataset

We employed the THINGS-fMRI dataset^24^. Every participant was presented with 8,640 natural images of 720 living and non-living objects (see Fig. 1b). Object categories and images were sampled from the THINGS database^22^. All analyses were computed using single-trial beta estimates created in a manner similar to GLMsingle^41,42^. Detailed procedures can be found in Hebart et al.^24^. For the analyses, we averaged the data across image examples of the 720 object categories and applied a spatial smoothing kernel of FWHM = 4 mm. For visualization of flattened visual cortex, we used the Python module pycortex^43^.

#### Vector-of-ROIs analysis

To create a set of sequentially, partially overlapping ROIs across OTC (Fig. 1d), we adapted the vector-of-ROIs approach described in Konkle and Caramazza^8^ to facilitate direct comparison with their findings. In the original implementation, the cortical trajectory was anchored in parahippocampal cortex (PHC), fusiform gyrus (Fus), inferior temporal gyrus (ITG), lateral occipital cortex (LO), and transverse occipital sulcus (TOS). These anatomical ROIs were not directly available in the THINGS-fMRI dataset. We therefore defined the trajectory using five available category-selective ROIs that spatially overlap with, or are close to, the regions used in the original approach.

Specifically, we chose the coordinates located in the centroids of the functionally defined medial place area (MPA), parahippocampal place area (PPA), fusiform face area (FFA), extrastriate body area (EBA), and occipital place area (OPA). Details on the creation of functional ROIs can be drawn from Hebart et al.^24^. We adjusted the anchor coordinate within EBA due to the relatively large size of the functional EBA-ROI, which also encompasses the middle temporal area (MT). As a result, much of the category-selective activation is primarily concentrated in the posterior regions of EBA. To ensure that the spheres map through peak activations of EBA, we defined a path from the EBA centroid to the V1 centroid. Along this trajectory, we selected the coordinate within the EBA that exhibited the highest activation across all single-trial beta estimates for all object categories as the new EBA anchor.

On the surface of the cortex, we then defined additional, evenly spaced coordinates falling onto a spline connecting the five anchors, resulting in 31 coordinates in total. Finally, we drew a spherical ROI (*r* = 10 mm) around each coordinate and projected the resulting spheres into the volume space.

We visualized how animacy and size jointly organize object responses by computing the mean beta weights for large animals (*n* = 24), small animals (*n* = 24), large objects (*n* = 130), and small objects (*n* = 130). Objects only included categories with an animacy score of 2 or below on a scale from 1 (very inanimate) to 7 (very animate). We excluded human faces and body parts to match the stimulus set of the original studies. However, excluding human categories did not change the overall pattern of results.

#### Smoothness of property topographies

For each participant and property, we fit a univariate voxel-wise OLS regression to obtain property coefficient maps (Fig. 5). To quantify uncertainty in the smoothness estimates and construct a shuffled baseline, we generated 1,000 bootstrap samples by resampling the 720 object categories with replacement. For each bootstrap sample, we estimated a coefficient map for each property. As a control, we additionally generated shuffled coefficient maps by randomly permuting the property scores across the resampled objects while leaving the corresponding fMRI responses unchanged.

We then quantified the spatial smoothness of each property coefficient map or shuffled coefficient map using a neighborhood-based, distance-weighted total-variation metric on the surface mesh. To this end, we projected the volumetric coefficient maps to the subject’s fiducial surfaces using pycortex. Coefficient maps were z-scored per subject to compare patterns independently of overall effect magnitude. Geodesic neighborhoods were built by running Dijkstra’s algorithm on the mesh to obtain shortest-path distances *d*(*i,j*) between vertices. For a Gaussian kernel with bandwidth *σ* = 3 mm, we included neighbors within a practical cutoff of 3*σ* = 9 mm. Each neighbor pair (*i,j*) was assigned a spatial distance 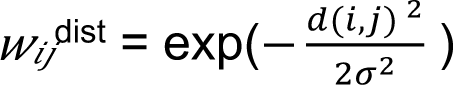 for *d*(*i,j*) ≤3*σ*, else 0.

The neighborhood-based, distance-weighted total-variation metric at scale *σ* was then computed as 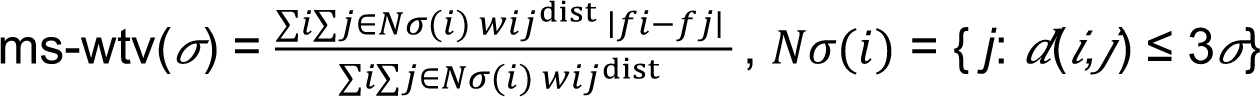, with *f*_i_ being the value of a given vertex *i* on a given property coefficient map. Lower values of ms-wtv(*σ*) indicate a smoother topographical organization.

Because vertices with low reliability can distort smoothness estimates, we computed a reliability-weighted variant that softly emphasizes coefficient differences between highly reliable vertices. For this, we used noise-ceiling maps provided in the THINGS-fMRI dataset^24^. Each participant’s noise-ceiling (nc) volume was projected onto the brain surface mesh, Gaussian-smoothed along the surface (FWHM = 6 mm), and min-max normalized to [0, 1]. For each neighbor pair (*i*,*j*), we defined a reliability weight reflecting the mean reliability value of a given vertex pair *w_ij_*^rel^ = max(mean(nc*_i_*, nc*_j_*), 0).

The distance- and reliability-weighted total-variation metric was then computed as: ms-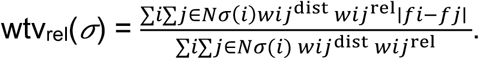

#### Hierarchical partitioning

To reduce the complexity of analyses, we combined the closely related properties “holdability” and “graspability” into “manipulability.” We excluded “manmadeness” due to its high correlation and conceptual overlap with “naturalness”, resulting in fifteen properties. As the object properties were still highly correlated, we opted for hierarchical partitioning to provide a more nuanced and fair assessment of each property’s independent contribution^26^. We constructed 2^15^ − 1 (= 32,767) ordinary least squares regression models to capture every possible combination of predictors, ranging from simple univariate regressions to a complete model incorporating all fifteen properties.

For cross-validation, we trained the models on 80% of the data and evaluated them on a separate 20% test set. We repeated this process over ten iterations of randomized train-test splits and averaged the performance results. For each property, we compared every model that included the property to its corresponding model with the same combination of predictors except that property. The property’s incremental variance was computed as the average *R*² difference across all such matched model pairs. The resulting voxel-wise *R*² estimates for each property were noise-normalized by dividing them by the noise ceiling of each voxel. Voxels with a noise ceiling below 5% were set to *R*² = 0.

### Statistics and reproducibility

All analyses were performed using Python 3.13. Unless stated otherwise, analyses were conducted at the level of object categories rather than individual object images. Sample sizes were determined by the available data in the THINGS-fMRI^24^ and THINGSplus^23^ datasets. Exact sample sizes, exclusion criteria, and definitions of sample sizes are reported in the corresponding Methods sections.

Because THINGS-fMRI is a densely sampled dataset, analyses were conducted separately for each participant, allowing consistency of results to be assessed across participants. For the smoothness metric and vector-of-ROIs analysis, participant-level results were subsequently averaged across participants. Reproducibility of model-based estimates, including explained variance in univariate encoding models and independent property contributions in the hierarchical partitioning analysis, was further assessed using ten randomized train-test splits.

## Acknowledgments

This work was supported by a Max Planck Research Group Grant (M.TN.A.NEPF0009) awarded to MNH and the ERC Starting Grant COREDIM (ERC-StG-2021-101039712) awarded to MNH.

## Data availability

This study used an existing publicly available fMRI dataset provided by the THINGS-data project^24^ available at https://openneuro.org/datasets/ds004192. Image stimuli and object property ratings were obtained from the THINGS/THINGSplus^22,23^ databases, with additional property ratings newly collected for the present study. Both the previously available and newly collected stimuli and property ratings are available at https://osf.io/jum2f. The data underlying the figures of this study are publicly available on

Figshare at 10.6084/m9.figshare.33395338.

## Code availability

Code used to perform the main analyses reported in this study will be made publicly available via GitHub upon publication at https://github.com/LauraStoinski/anisize.git.

## Author contributions

Conceptualization and Methodology: M.N.H., T.K., and L.M.S. Formal analysis and Visualization: L.M.S. Funding acquisition: M.N.H. Supervision: M.N.H. and T.K. Writing original draft: L.M.S. Review and editing: M.N.H., T.K. and L.M.S.

