## Supplemental Information for "Principles of coarse-scale functional organization in occipitotemporal cortex"

Laura M. Stoinski

This PDF file includes:

Supplementary Methods and Results

Supplementary Figure 1

Supplementary Figure 2

Supplementary Figure 3

Supplementary Figure 4

Supplementary Table 1

Supplementary Table 2

Supplementary Table 3

Supplementary References

### Supplementary Methods and Results

*Robustness of full factorial organization of animacy and size to stimulus effects*

We conducted various exploratory analyses to evaluate the robustness of our animacy and size topographies (Supplementary Fig. 1). First, we confirmed that the additional size preference regions were also visible in the volumetric data, ensuring they were not a result of erroneous surface mapping. Second, we ensured that the size topography remained consistent when neglecting the continuity of our size measure, only comparing sensitivities to the 25% largest and the 25% smallest categories, similar to how it was done in the original work[1].

Further, we investigated the influence of category diversity. The original study[1] did not include images of humans or food. In light of the strong response preferences of FFA and EBA to human faces and bodies[2,3] and the recently proposed food-selective region between FFA and PPA[4,5], we tested the influence of these categories on our results. However, excluding images of food, human faces and bodies, or even repeating the analysis only with objects included in the original study ($n$ = 101), did not alter the more fine-grained size partition. Animate and inanimate categories covered largely overlapping size ranges with only ten objects falling outside the range spanned by animals, which extended from "ant" to "whale". However, compared to the original work[1], our categories varied more strongly in their size, ranging from a small grain of sand up to an aircraft carrier. To investigate the impact of size range, we repeated the analysis excluding extremely large (size > 320) and extremely small (size < 140) categories (excluded $n$ = 84). We also compared response to only the 25% largest and smallest categories. Yet, we observed a similar pattern of responses, indicating that this effect is not solely due to the wider range and continuity of our size ratings.

We additionally explored whether characteristics other than size might explain the additional size regions in our data. Size was only weakly correlated with the eccentricity ($r$ = 0.02, $p$ = 0.16) or the size at which the objects are displayed in the images ($r$ = 0.06, $p$ < 0.001). We compared animacy and size ratings with image-wise estimates of other visual dimensions, as determined from human similarity judgments of the THINGS images[6–8]. Animacy and size were only weakly correlated with color-related dimensions (all $|r|$ < 0.16). However, size correlated with the "coarse-scale pattern/many things" dimension ($r$ = -0.49, $p$ < 0.001), as smaller categories were more frequently portrayed collectively in pattern-like arrangements (e.g., a pile of lentils). However, excluding fine-grained pattern-like images (i.e., patternlikeness > 0.6, excluded $n = 354$) before averaging the responses across images showing the same concept did not alter the observed animacy and size response profiles.

Finally, the original studies[1,9] may have overlooked finer size partitions as they assessed size topography in group averages rather than individual participants, as done here, potentially obscuring these details in the spatially normalized and smoothed results. However, even after testing the effects of averaging and applying a broad smoothing kernel (FWHM = 6–8 mm), we continued to observe a finer size partition.

*Smoothness of property topographies after controlling for animacy*

To explore whether the smoothness of properties correlated with animacy merely reflects variance shared with animacy, we regressed animacy-related variance out of both the property scores and the fMRI responses and then computed regression slope maps for the residualized object properties. Smoothness was quantified using the same procedure described in the Smoothness of property topographies analysis in the Methods section of the main manuscript, with the exception that we used 10 bootstrap iterations instead of 1,000 to reduce computational demands for this exploratory analysis. Supplementary Fig. 2a shows the resulting topographical maps after controlling for animacy, and Supplementary Fig. 2b shows the corresponding smoothness scores. Even after removing variance associated with animacy, many properties continued to produce broad and spatially smooth maps, suggesting that their topographical smoothness cannot be explained solely by their correlation with animacy.

*Participant-wise differences in hierarchical partitioning results*

To assess the consistency of property effects across participants, we plotted the results shown in Fig. 7 of the main manuscript separately for each participant. Supplementary Fig. 3a shows the variance explained by each property, estimated using univariate OLS regression models. Supplementary Fig. 3b shows the incremental variance explained by each property, estimated using hierarchical partitioning.

Overall, participant-wise patterns were broadly consistent with the group average. Moveability and humanness consistently remained the properties explaining most incremental variance in FFA and EBA. In other regions, the ranking of the properties explaining most incremental variance varied slightly more across participants. In PPA, size, naturalness and hardness explained most incremental variance in S1; the ability to be moved and size in S2; and the ability to be moved and size in S3. In the previously proposed small-object region between FFA and EBA[1], size and heaviness explained the most incremental variance in S1 and S3, whereas moveability explained most incremental variance in S2. Together, these results demonstrate that the pronounced effects of moveability and humanness in FFA and EBA, as well as the more balanced contribution of multiple properties to explaining responses in other regions, are consistent across participants.

*Robustness of hierarchical partitioning results to modulations of the predictor set*

Because many of the investigated properties were correlated with animacy or size, we further examined whether excluding highly correlated properties affected the hierarchical partitioning results (for correlations, see Fig. 4a of the main manuscript). We first repeated the analysis without naturalness and heaviness, which were strongly correlated with animacy or size, respectively. We then additionally excluded agency because of its high correlation with animacy.

As shown in Supplementary Fig. 4, both analyses yielded highly similar results. As expected, including fewer predictors slightly increased the incremental variance explained by the remaining predictors. This was particularly the case for animacy and size in PPA and OPA. As in the full analysis (Fig. 7c of the main manuscript), moveability and humanness explained more incremental variance than animacy in FFA and EBA, and size, animacy, and moveability explained the most incremental variance in PPA. In OPA, however, animacy explained the most incremental variance after excluding correlated predictors, whereas the full analysis showed a more balanced contribution of animacy, size, and the ability to be moved. Overall, these results show that the strong effects of moveability and humanness beyond animacy are robust and do not simply reflect a bias of including multiple animacy-related properties in the full property set.

*Exploration of objects eliciting the highest and lowest responses within spherical ROIs.*

To identify the object concepts eliciting the highest and lowest responses in each spherical ROI, we averaged the single-trial beta estimates provided by the THINGS-fMRI dataset[7] across the 12 images depicting each object concept, and then averaged these object-wise responses across participants. Supplementary Table 1 shows the 15 objects eliciting the highest responses and the 15 objects eliciting the lowest responses within a subset of ROIs overlapping with animacy- or size-preference regions identified in the present study (see Fig. 2 of the main manuscript).

The resulting profiles broadly matched known animacy-, size-, and category-selective organization in occipitotemporal cortex. ROIs previously related to large-object and scene preferences, including PPA, MPA, and OPA[1,10], showed stronger responses to large inanimate objects, such as subway, submarine, and carousel, and weaker responses to animate or natural categories, such as bear and rabbit, as well as smaller inanimate objects, such as cup and blowtorch. In animacy-, face-, and body-selective ROIs, including FFA and EBA[1–3], the strongest responses were elicited by animate concepts, body parts, people, animals, and categories often depicted with bodies, such as clothing, whereas the weakest responses were predominantly elicited by inanimate objects.

However, there were exceptions to this pattern. For example, small inanimate objects such as credit cards elicited relatively strong responses in OPA, and body-related concepts such as hip also appeared among higher-ranking concepts. Thus, the ROI response profiles broadly followed known animacy-, size-, and category-preference gradients, while also showing concept-specific deviations not captured by broad labels such as animate, inanimate, large, or small.

*Exploration of objects with lowest or highest prediction errors within spherical ROIs based on property encoding models*

To explore which objects contributed to the larger shares of explained variance for moveability and humanness compared to animacy in FFA and EBA (Fig. 7b and c of the main manuscript), we examined object-wise prediction errors from the corresponding univariate encoding models of these properties. We additionally included the prediction errors of the size and agency encoding models, as these models also explained substantial variance in these regions.

Supplementary Tables 2 and 3 list the objects with the 15 lowest absolute, the 15 highest positive and the 15 highest negative residuals in FFA and EBA, respectively. Residuals were defined as measured minus predicted fMRI responses. Low absolute residuals thus indicate objects whose fMRI responses were well predicted by the model. High positive residuals indicate objects for which the model underestimated the measured response, whereas negative residuals indicate objects for which the model overestimated the measured response.

In both FFA (Supplementary Table 2) and EBA (Supplementary Table 3), objects with the largest positive residuals included several clothing categories and body parts. These categories may receive relatively low animacy, moveability, agency, or humanness ratings despite depicting or strongly resembling aspects of the human body. For the animacy model, responses to plants, which received relatively high animacy ratings among non-animal categories, were frequently overestimated. For the moveability model, vehicles were among the objects whose responses were overestimated. Thus, the residual patterns provide additional insight into which object categories are not well captured by individual predictors and suggest that the higher contribution of moveability than animacy in FFA and EBA is unlikely to simply reflect a general preference for both animals and highly moveable inanimate objects such as vehicles.

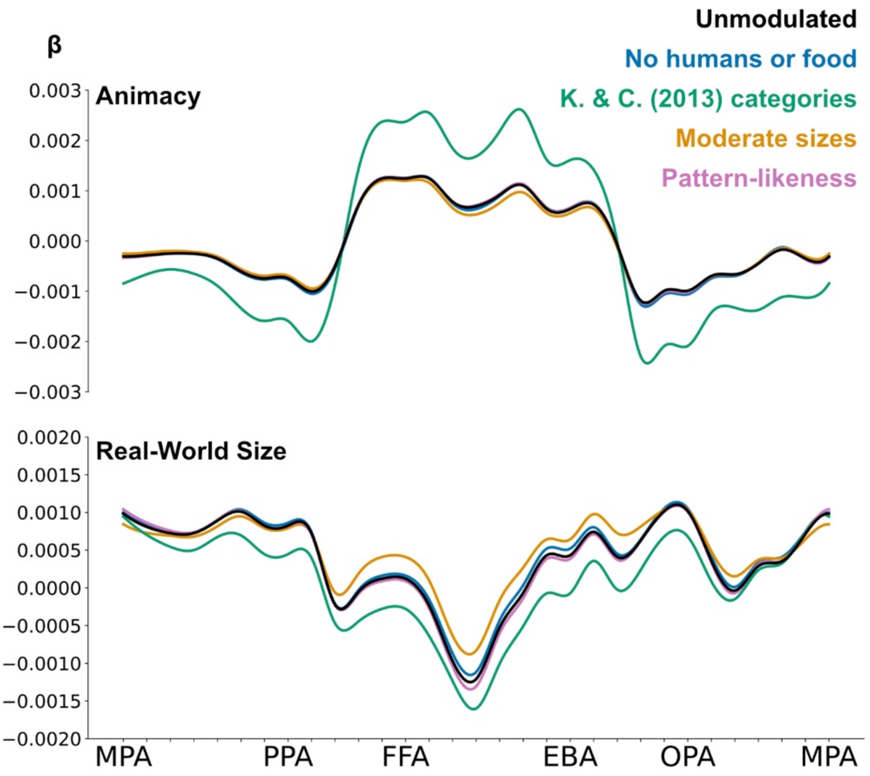


**Supplementary Fig 1. Robustness of animacy and size response profiles to exploratory control analyses.** To assess the stability of the observed animacy and real-world size organization, we recomputed response profiles in high-level visual cortex after systematically modifying the stimulus set. Profiles are averaged across the three participants. Control analyses were designed to approximate the stimulus set of the original study and to test potential stimulus-driven confounds. Specifically, we excluded images containing humans or food, restricted analyses to the object categories used in Konkle & Caramazza[1], removed categories with extremely small or large real-world sizes, and excluded images with highly pattern-like structure. Across these modulations, the finer-grained size response patterns remained similar, indicating that the reported organization is unlikely to be driven by these stimulus factors.

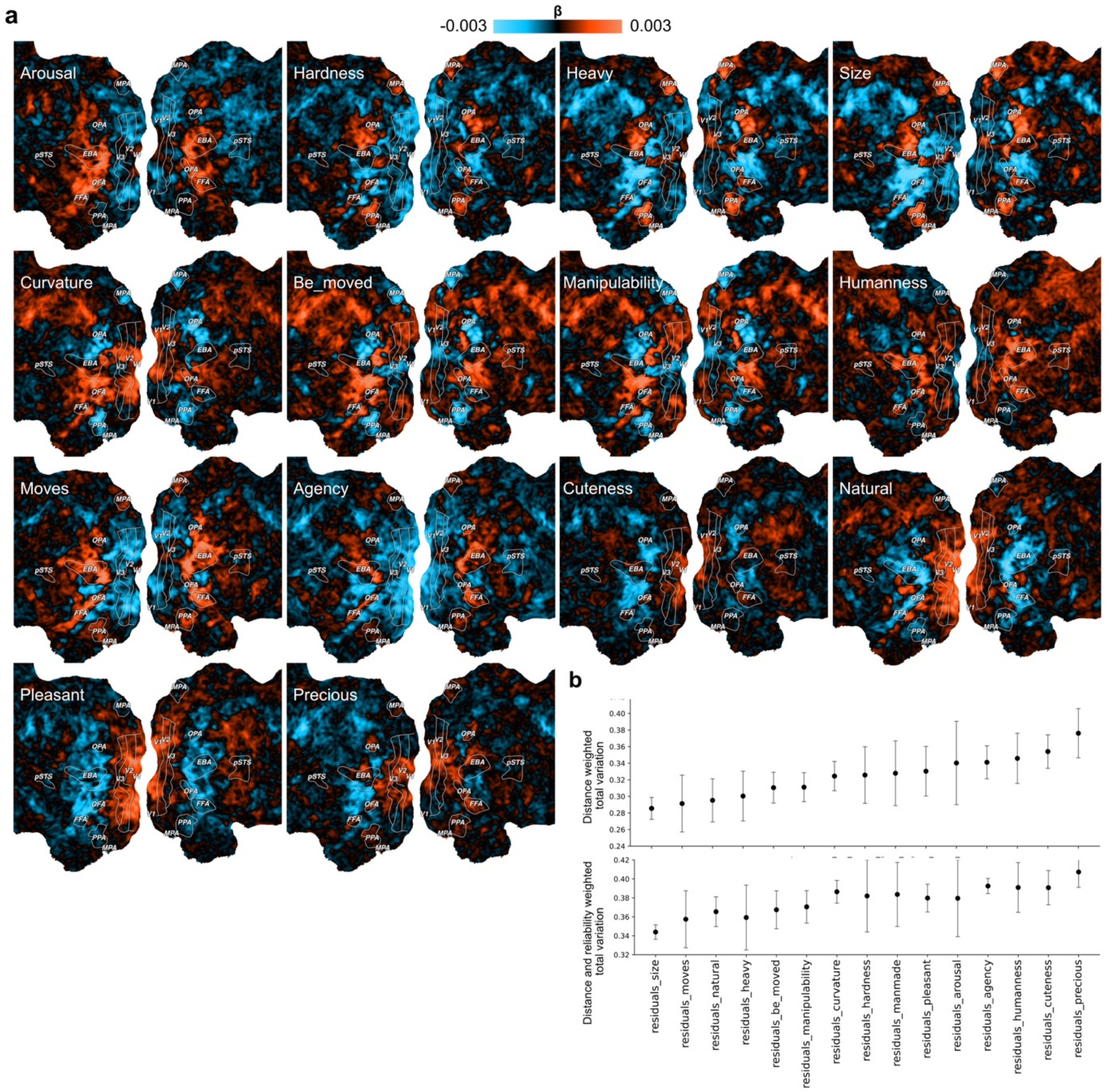


**Supplementary Fig 2. Smoothness of topographic property maps after controlling for animacy. a.** Topographic maps of object properties after regressing animacy-related variance out of property scores and fMRI responses. **b.** Corresponding smoothness scores for the residualized property topographies. Many properties continued to exhibit broad and spatially smooth topographies after controlling for animacy, indicating that their smoothness cannot be explained solely by variance shared with animacy.

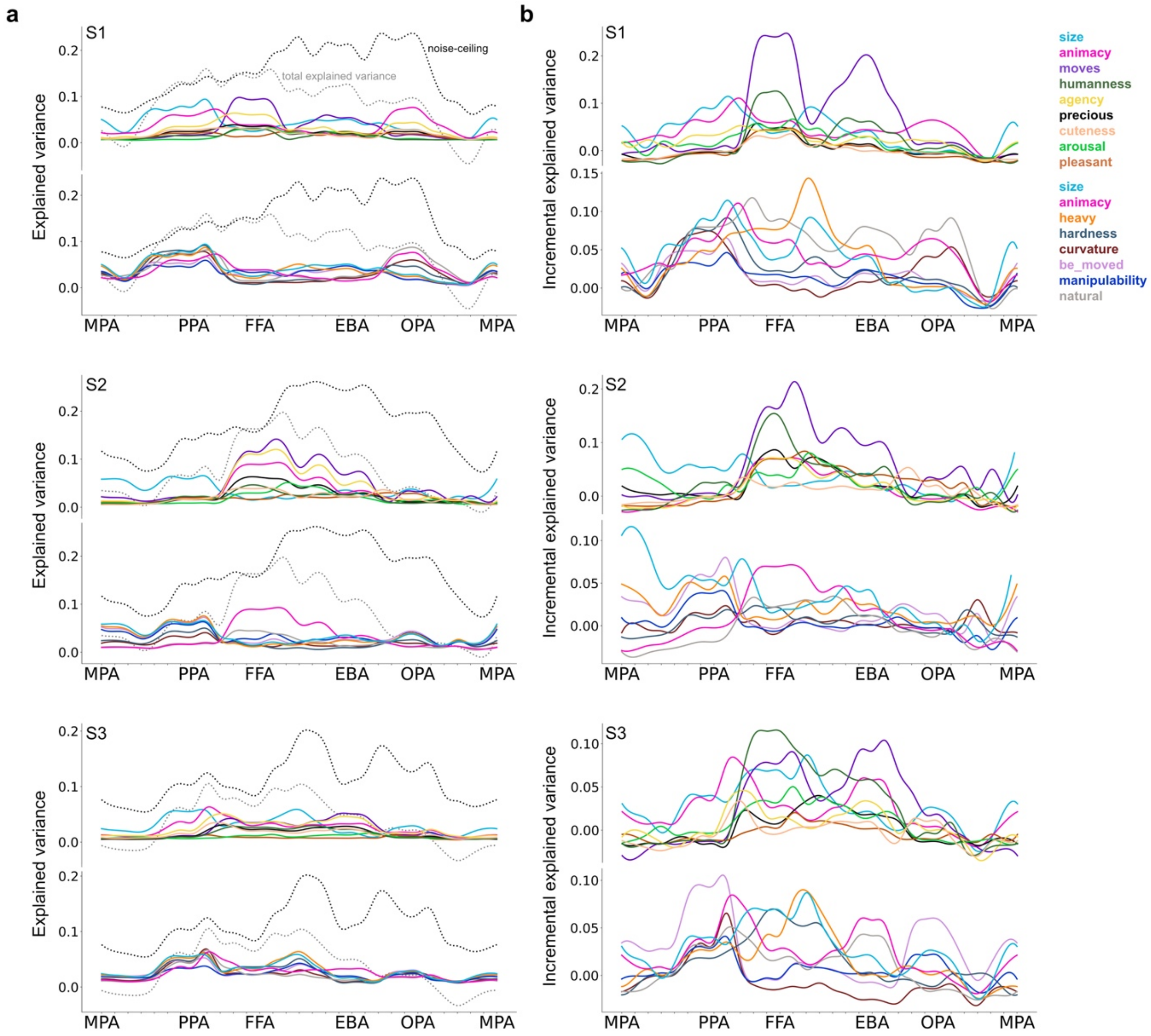


**Supplementary Fig 3. Participant-wise explained variance by object property across high-level visual cortex. a.** Noise ceiling of the fMRI data, and total variance explained by univariate models for each property regressor and full model including all properties (cross validated). Across participants, moveability, agency, animacy, and size explained the largest shares of variance overall. **b.** Incremental variance explained by each property, estimated using hierarchical partitioning (cross-validated and noise-normalized). Across participants, moveability and humanness consistently explained most incremental variance in FFA and EBA. In other regions, incremental variance was more evenly distributed across properties, and the ranking of properties explaining the most incremental variance varied slightly more across participants.

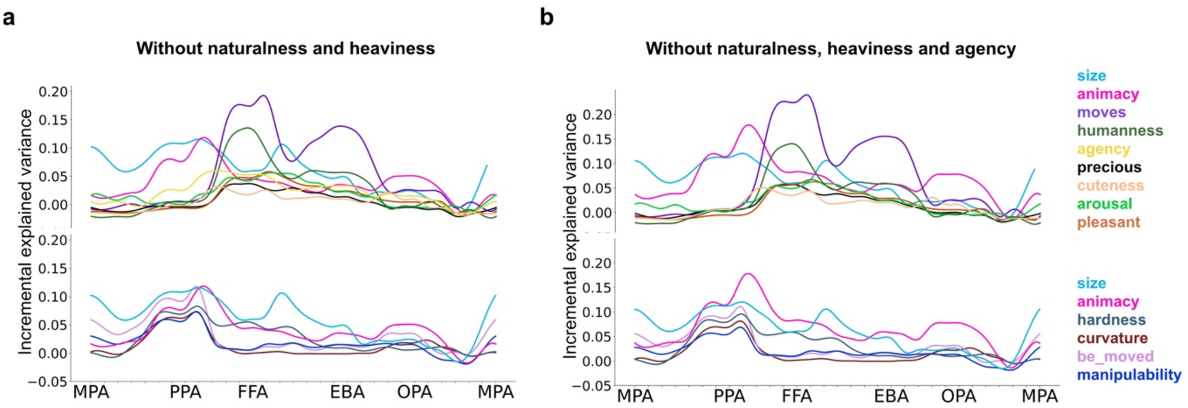


**Supplementary Fig 4. Effects of modifying the set of predictors on the hierarchical partitioning analysis. a.** Hierarchical partitioning results after excluding naturalness and heaviness. The resulting pattern was highly similar to the full analysis, with moveability and humanness explaining most incremental variance in FFA and EBA. **b.** Hierarchical partitioning results after additionally excluding agency. The overall pattern remained similar to the full analysis, with moveability and humanness explaining most incremental variance in FFA and EBA.

**Supplementary Table 1. Object concepts eliciting the highest and lowest responses in a selection of spherical ROIs.**

| Sphere | 7 (PPA) | 8 | 12 (FFA) | 15 | 19 (EBA) | 24 (OPA) | 30 (MPA) |
|---|---|---|---|---|---|---|---|
| Highest response in ROI | subway | waterfountain | chest1 | hip | hip | submarine | submarine |
| | desk | bumper | hip | chest1 | pogo_stick | hammock | bed |
| | tent | desk | girl | lingerie | footbath | pothole | hammock |
| | bumper | subway | woman | helmet | chest1 | pogo_stick | waterfountain |
| | submarine | pothole | pogo_stick | garter | leggings | airbag | treadmill |
| | waterfountain | submarine | face | tadpole | iceskate | waterfountain | carousel |
| | pothole | tent | shower_cap | snorkel | snorkel | footbath | pogo_stick |
| | railing | mosquito_net | leggings | leggings | footrest | tent | clay |
| | stalagmite | footbath | garter | octopus | anklet | credit_card | tablecloth |
| | mosquito_net | hammock | hamster | tattoo | dog | desk | eyepiece |
| | hammock | guillotine | snorkel | wooden_leg | rollerblade | helmet | banana |
| | sodafountain | bed | monkey | anklet | kimono | hip | subway |
| | solar_panel | meat_grinder | dog | pacifier | thumbtack | guillotine | manatee |
| | seesaw | dishwasher | kimono | retainer | hamster | computer_screen | snowboard |
| | meat_grinder | wooden_leg | doll | prism | snake | subway | doormat |
| Lowest response in ROI | pumpkin | bungee | kettle | cage | radiator | lettuce | bag |
| | bear | kettle | radiator | beaker | croissant | bag | cinnamon |
| | squirrel | squirrel | cup | fire | altar | peanut | rabbit |
| | potato | peanut | lime | dollhouse | fern | steak | bandage |
| | lime | sheep | fence | christmastree | manhole | tray | uniform |
| | cup | rabbit | parsley | tree | fence | cookie | nacho |
| | stethoscope | cup | cherry | speaker | jar | lime | record |
| | macadamia | stethoscope | speaker | poster | beaker | possum | hammer |
| | cashew | uniform | fern | cherry | scaffolding | croissant | cufflink |
| | croissant | potato | scaffolding | altar | speaker | sheep | stethoscope |
| | cherry | hand | cage | fence | cage | macadamia | blowtorch |
| | hand | cherry | altar | parsley | couch | parsley | elbow |
| | rhinoceros | croissant | towel_rack | scaffolding | cherry | breadstick | butterfly |
| | kettle | rhinoceros | couch | couch | parsley | bamboo | mousse |
| | parsley | parsley | bamboo | bamboo | bamboo | cherry | tray |

**Supplementary Table 2. Object concepts with the lowest and highest residuals in spherical ROI 12 (FFA) for selected property-based encoding models**

| Model: | Animacy | Size | Moves | Humanness | Agency |
|---|---|---|---|---|---|
| Lowest absolute residuals | burner | corn | cauliflower | subway | pocketknife |
| | chocolate | egg | blanket | velcro | penguin |
| | mosquito | rocking_chair | lasagna | egg | syringe |
| | sleeping_bag | wheelbarrow | possum | eyepiece | joystick |
| | bee | snowplow | icicle | granite | playing_card |
| | groundhog | space_shuttle | bear | thimble | ping-pong_table |
| | anchor | mop | celery | broom | porcupine |
| | donut | pie | crutch | corn | hedgehog |
| | birdcage | missile | chalk | camcorder | frog |
| | nest | ping-pong_table | chalice | quill | doormat |
| | hovercraft | bird | candelabra | crayon | pecan |
| | popsicle | pothole | hanger | jersey | meatball |
| | zebra | icicle | bed | needle | corkscrew |
| | trap | crayon | soda_fountain | hatbox | snowplow |
| | mosquito_net | oyster | fish | donkey | treadmill |
| Highest positive residuals | hip | chest1 | chest1 | chest1 | hip |
| | chest1 | hip | hip | pogo_stick | chest1 |
| | pogo_stick | girl | shower_cap | hip | pogo_stick |
| | shower_cap | woman | garter | otter | shower_cap |
| | leggings | pogo_stick | leggings | snorkel | leggings |
| | garter | shower_cap | snorkel | wolf | snorkel |
| | snorkel | leggings | doll | shower_cap | hood |
| | doll | garter | pogo_stick | dog | kimono |
| | girl | hamster | kimono | girl | garter |
| | kimono | snorkel | girl | lobster | girl |
| | woman | monkey | tattoo | tadpole | anklet |
| | hood | dog | helmet | hamster | woman |
| | robot | doll | woman | platypus | doll |
| | helmet | kimono | hood | woman | chin |
| | tattoo | wolf | footbath | warthog | footbath |
| Highest negative residuals | bamboo | bamboo | bamboo | bamboo | bamboo |
| | parsley | couch | bus | couch | wheat |
| | wheat | altar | couch | cherry | parsley |
| | cherry | scaffolding | tractor | scaffolding | cherry |
| | couch | cage | fire | kettle | couch |
| | lime | speaker | wheat | altar | altar |
| | hedge | parsley | golf_cart | lime | bus |
| | pumpkin | cherry | aircraft_carrier | speaker | fire |
| | altar | fence | parsley | parsley | speaker |
| | scaffolding | lime | cage | cage | giraffe |
| | cage | radiator | giraffe | fence | cage |
| | grape | wheat | scaffolding | hairpin | lime |
| | flower | tray | wheelchair | bus | radiator |
| | peanut | kettle | mast | dresser | hedge |
| | speaker | bus | bike | croissant | scaffolding |

**Supplementary Table 3. Object concepts with the lowest and highest residuals in spherical ROI 19 (EBA) for selected property-based encoding models**

| Model: | Animacy | Size | Moves | Humanness | Agency |
|---|---|---|---|---|---|
| Lowest absolute residuals | spatula | goalpost | filing_cabinet | undershirt | crown |
| | possum | inhaler | book | comb | life_jacket |
| | inhaler | pecan | papaya | pear | hanger |
| | life_jacket | walker1 | sword | paint | lemon |
| | rocking_chair | cow | toothpick | blazer | grate |
| | garbage | sushi | zucchini | crib | spatula |
| | blender | polaroid | bench | key | milk |
| | polaroid | fish | ferris_wheel | cellphone | ruler |
| | grate | blazer | dishwasher | boy | skunk |
| | hanger | bulldozer | ruler | blind | boxing_gloves |
| | tomato | drum | chicken_wire | egg | donut |
| | antenna | crayon | peeler | antenna | sushi |
| | bib | net | leek | tiara | spider |
| | milk | airplane | pan | stair | lollipop |
| | projector | onion | football_helmet | garbage | raft |
| Highest positive residuals | pogo_stick | hip | pogo_stick | pogo_stick | pogo_stick |
| | hip | pogo_stick | hip | hip | hip |
| | footbath | footbath | footbath | footbath | footbath |
| | leggings | leggings | leggings | snorkel | iceskate |
| | iceskate | chest1 | snorkel | footrest | footrest |
| | snorkel | snorkel | footrest | iceskate | anklet |
| | footrest | iceskate | anklet | thumbtack | leggings |
| | anklet | anklet | iceskate | snake | snorkel |
| | chest1 | footrest | thumbtack | dog | chest1 |
| | rollerblade | thumbtack | kimono | leggings | rollerblade |
| | thumbtack | floss | floss | anklet | thumbtack |
| | kimono | dog | credit_card | bow2 | kimono |
| | floss | hamster | chest1 | chest1 | bow2 |
| | bow2 | rollerblade | brace | dolphin | boot |
| | boot | snake | boot | rollerblade | wooden_leg |
| Highest negative residuals | bamboo | bamboo | bamboo | bamboo | bamboo |
| | parsley | couch | parsley | cherry | parsley |
| | cherry | cage | cage | parsley | cherry |
| | couch | parsley | cherry | couch | speaker |
| | cage | speaker | couch | speaker | cage |
| | speaker | cherry | speaker | cage | couch |
| | scaffolding | scaffolding | tractor | scaffolding | beaker |
| | beaker | fence | bus | croissant | scaffolding |
| | jar | beaker | scaffolding | jar | jar |
| | asparagus | jar | beaker | nacho | altar |
| | fence | altar | rhinoceros | fence | radiator |
| | croissant | radiator | golf_cart | beaker | fence |
| | tree_trunk | dollhouse | aircraft_carrier | bottle | asparagus |
| | altar | croissant | banner | altar | croissant |
| | potato | rug | jar | dresser | bottle |
